# Cryopreserved cGMP-compliant human pluripotent stem cells-derived immature hepatic progenitors rescue mice from acute liver failure

**DOI:** 10.1101/2022.09.26.509491

**Authors:** Malika Gantier, Raphael Rispal, Angélique Fourrier, Séverine Menoret, Frédéric Delbos, Sarah Renault, Anne-Sophie Gary, Ignacio Anegon, Tuan Huy Nguyen

## Abstract

Liver transplantation remains the only curative treatment for end-stage liver diseases. Unfortunately, there is a drastic organ donor shortage. Hepatocyte transplantation emerged as a viable alternative to liver transplantation. In light of their unique expansion capabilities and their potency to be driven towards a chosen cell fate, pluripotent stem cells (PSC) are extensively studied as an unlimited cell source of hepatocytes for cell therapy. It has been previously shown that freshly prepared hepatocyte-like cells can cure mice from acute and chronic liver failures and restore liver functions. In this study, we generated human PSC-derived immature hepatic progenitors (GStemHep) using current good manufacturing practice (cGMP) compliant conditions from PSC amplification, hepatic differentiation to cell cryopreservation. These GStemHep cells present an immature hepatic phenotype (alpha-fetoprotein positive, albumin negative), secrete hepatocyte growth factor (HGF) and do not express MHC type I or II. The therapeutic potential of GStemHep was assessed in two clinically relevant models of acute liver failure. A single dose of thawed GStemHep rescue mice from sudden death caused by acetaminophen and thioacetamide-induced acute live failure, both in immunodeficient and immunocompetent animals in absence of immunosuppression. The mode of action was studied by several analytical methods including unbiased proteomic analyses. The swiftness of the therapeutic effect suggests a paracrine mechanism of action of GStemHep leading to a rapid reduction of inflammation and a rapid cytoprotective effect. Therapeutic biological effects were observed as soon as 3 hours post-cell transplantation with reduction in serum transaminases and in liver necrosis. Mode of action of GStemHep relies on alleviation of inhibition factors of liver regeneration, increase in proliferation-promoting factors and decrease liver inflammation. In conclusion, we generated cGMP-compliant human PSC-derived immature hepatic progenitors that were highly effective in treating acute liver failure. This is also the first report highlighting that human allogeneic cells could be used as cryopreserved cells and in absence of immunosuppression for a human PSC-based regenerative medicine of acute liver injuries.

## Introduction

Acute liver failure (ALF) is a rare and severe liver disease that has a high mortality and affects many organ systems. It is characterized by a rapid and massive deterioration of liver functions, which results in hepatic encephalopathy and coagulopathy within 8 weeks of the first symptoms in individuals with no preexisting liver disease (Bernal et al., 2010; Lefkowitch, 2016). The main causes of this critical pathology are drug-induced liver injury, intoxications, viral hepatitis, autoimmune liver disease and shock or hypoperfusion (Lefkowitch, 2016; Polson and Lee, 2005).

So far, orthotopic liver transplantation (OLT) is the only curative treatment for severe forms of ALF at end-stage. Unfortunately, because of a drastic shortage of organ donors many patients are excluded from the liver waiting list with exclusion criteria such as cancer outside the liver, substance abuse, disabling psychiatric conditions or lack of inadequate insurance. In addition, this treatment requires a heavy surgery with higher mortality risks for elderly patients and/or patients with pre-existing cardiac illness and a lifelong immunosuppressive treatment with known side effects such as infections, renal toxicity and a higher risk of cancer (Hernández Vallejo et al., 2005).

Hepatocyte transplantation (HT) isolated from discarded livers for transplantation due to quality criteria has become an attractive alternative or a bridge to OLT (Dhawan, 2015; Dhawan et al., 2010; Hansel et al., 2014). This alternative has been tested in patients with ALF or inherited metabolic diseases such as Crigler-Najjar syndrome type 1, urea cycle defects or factor VII deficiency (Lee et al., 2018). HT has demonstrated positive results by improving patients’ liver functions with decreased blood bilirubin levels, spontaneous healing, or successful transition to liver transplantation. However, the results obtained from these studies could not define the exact conditions and population of patients for which this therapy would be beneficial (Iansante et al., 2018; Lee et al., 2018). This can be explained by the limited number of patients, the variable quality of the isolated hepatocytes, and the different protocols used between care centers. Moreover, current culture conditions failed to efficiently expand *in vitro* primary hepatocytes to a sufficient scale for the treatment of a large number of patients resulting in a continual need for multiple liver donors (Garnier et al., 2018; Hu et al., 2018; Huch et al., 2015). HT has paved the way to cell therapy as an alternative to OLT but a new source of cells, available of constant quality at any time and on a large scale, is needed.

Pluripotent stem cells (PSC) have emerged as the ideal cell source. Indeed, they have the capacity for unlimited self-renewal and to differentiate into any cell types of the body, including hepatocytes. Many teams have already developed protocols for the differentiation of PSCs into hepatocytes, so-called hepatocyte-like cells (HLC), which not have a fully mature hepatic phenotype as they still expressing alpha-fetoprotein (AFP), a marker of fetal hepatocytes (Blackford et al., 2019; Hannan et al., 2013; Luce et al., 2021; Rashidi et al., 2018; Schwartz et al., 2014). The therapeutic potential of HLCs has been demonstrated in ALF animal models using freshly prepared cells and non-compliant current good manufacturing practices (cGMP) conditions, e.g. due to the use of Matrigel as an extracellular culture matrix (Chen et al., 2020; Nagamoto et al., 2016; Pettinato et al., 2016, 2019; Tolosa et al., 2015). While these results are encouraging, HLC transplantation in humans presents serious challenges. Indeed, similarly to primary hepatocytes, these cells might be highly sensitive to cryopreservation and lost most of their therapeutic potential upon cell thawing (Stéphenne et al., 2007; Terry et al., 2010). However, it is very important to cryopreserve the cells to always guarantee the availability of the treatment at any time in the care centers. While cell cryopreservation being a key point for emergency treatment of patients with end-stage liver diseases, only freshly prepared HLCs were used for treating ALF animal models. We have previously shown that the cryopreserved PSC-derived hepatic cells can treat an inherited metabolic liver disease, but this is a pathological context where hepatic cells have time to engraft and to mature into fully differentiated hepatocytes (Fourrier et al., 2020). Moreover, the current production protocols must comply to cGMP to be used as medicine in the clinic. Few studies present cGMP compliant protocols and none have evaluated them for the treatment of animals modeling acute or chronic liver diseases using cGMP compliant HLCs (Graffmann et al., 2022). In addition, as previously shown in humans, there is a poor integration of transplanted cells in the host liver despite immunosuppression, suggesting a poor understanding of the mechanisms leading to failure of cell implantation and/or cell rejection as well as a lack of efficient ways to prevent it (Allen et al., 2008; Han et al., 2009; Olszewski et al., 2001).

Here, we describe the production, the cryopreservation and the characterization of human PSC-derived hepatic progenitors called GStemHep generated using cGMP-compliant protocols. These cells present an immature hepatic phenotype (positive for AFP, negative for albumin/ALB), secrete HGF and have a low immunogenic profile (no expression of CMH class I or II). We show the therapeutic effect of cryopreserved GStemHep *in vivo* in two different models of ALF, i.e. acetaminophen (APAP) overdose in immunodeficient mice (NOD/SCID) and thioacetamide (TAA) overdose in immunocompetent mice (C57Bl/6) in absence of immunosuppression. GStemHep survival after transplantation and mechanisms of action have been studied. We show that cryopreserved cGMP-compliant GStemHep rescue mice from ALF with a better understanding of the mechanism of action and its kinetics following cell transplantation. We have overcome the current challenges in liver cell therapy with an unlimited cell production from a same starting cell raw material (human PSC) to avoid variable quality of cell therapy products due to multiple donor sourcing (inherent inter-individual variability), cGMP compliant therapeutic cells (PSC amplification and hepatic differentiation) and cryopreserved cells for immediate clinical availably.

## Materials and Methods

### Cell culture

Human PSC were provided by ESI BIO (Alameda, USA) and derived under cGMP conditions on human fibroblast feeder layers and cell lines are available in research and GMP-grade formats (Crook et al., 2007). They were adapted and then cultured in feeder-free conditions on culture dishes coated with the Vitronectin Recombinant Human Protein (ThermoFisher) in mTeSR1™ medium (StemCell Technologies) at 37°C in a 5% CO2 incubator with daily media changes and were passaged every 4-5 days using TrypLE^TM^ (ThermoFisher), following manufacturer’s recommendations. After each passage, they were cultured during 24 hours in the presence of 10 μM of Rock inhibitor Y-27632 (StemCell Technologies).

### In Vitro hepatic differentiation and GStemHep culture

PSCs were plated onto iMatrix-511 (Amsbio) in mTeSR1™ with 10 μM of Rock inhibitor Y-27632, following manufacturer’s recommendations. After 48 hours, PSC maintenance medium was replaced by RPMI-1640 supplemented with B27 serum-free supplement (Life Technologies) to start differentiation (Day 0). Medium was changed daily thereafter following a 3 stages differentiation plan. During the first two days of definitive endoderm induction, cells were cultured in the presence of 3 μM CHIR-99021 (Stem Cell Technologies). Then cells were cultured for 1 day without cytokines. To induce hepatic cell specification, medium was supplemented with 10 ng/mL fibroblast growth factor 10 (FGF-10) (Miltenyi Biotec) and 10 ng/mL bone morphogenetic protein 4 (BMP-4) (R&D System) for five days. Finally, cells were cultured in presence of 3 μM CHIR-99021 and 20 ng/mL HGF (Miltenyi Biotec) for two days to obtain the GStemHep. After differentiation, cells were harvested and frozen at −150°C in CryoStor® CS10 cell freezing medium (StemCell Technologies). Cell count and viability were evaluated using NC-200 NucleoCounter (Chemometec). For the plating experiments, GStemHep were thawed and seeded onto iMatrix-511 coated-culture dishes in RPMI-1640 supplemented with B27, 10% FBS, 10 μM Y-27632, 3 μM CHIR-99021 and 20 ng/mL HGF.

### Animals

Male NOD/SCID mice and C57Bl/6 mice were obtained from Janvier Labs (Saint-Berthevin, France) and maintained in the animal facility on 12h light/dark cycles with food and water *ad libitum.* All animal care and procedures performed in this study were approved by the Animal Experimentation Ethics Committee of Pays de la Loire region, France, in accordance with the guidelines from the French National Research Council for the Care and Use of Laboratory Animals (Authorisation Numbers: Apafis 21417)

### APAP-induced ALF model

NOD/SCID mice (6 weeks-old) were treated with an intraperitoneal (ip) injection of 650 mg APAP/kg (Sigma-Aldrich) to induce ALF. Three hours later, animals received an intrasplenic injection of 1×10^6^ cryopreserved GStemHep, which were thawed, pelleted by a low-speed centrifugation and resuspended in RPMI/B27 medium. The control mice received APAP but no cells treatment. Mice survival was followed over 10 days or mice were euthanized at 6h, 9h and 24h post-APAP.

### TAA-induced ALF model

C57Bl/6 mice (6 weeks-old) were treated with an ip injection of 1500 mg TAA/kg (Sigma-Aldrich) to induce ALF. Twenty-four hours later, animals received an intrasplenic injection of 1×10^6^ cryopreserved GStemHep, which were thawed and directly used for cell injection. The control mice received TAA but no cells treatment. Mice survival was followed over 7 days or mice were euthanized at 24h.

### RNA extraction and real-time quantitative Polymerase Chain Reaction (RT-qPCR)

Total mRNA was extracted from cell culture using the RNeasy Mini kit (Qiagen) following the manufacturer’s recommendations. Total RNA from frozen liver was extracted using Trizol (Life Technologies) and chloroform (VWR) after mechanical disruption of the tissue. Then, the RNeasy Mini kit was used as for cell culture mRNA. RT-qPCR was performed with 5 ng of mRNA, with a one-step RT-PCR kit using Taqman® technology (AgPath-IDTM One-Step RT-PCR, Life Technologies) and using the Applied Biosystems ViiA 7 Real-Time PCR System and the appropriate primers for Taqman assays (Life Technologies; Supplementary Table 1). The relative gene expression was calculated using the 2-ΔΔCt quantification method after normalization to GAPDH values and expressed as fold of levels found in undifferentiated hPSCs cells for cultured cell or in WT mice group for liver analysis.

### Flow Cytometry

Cryopreserved GStemHep were thawed, centrifuged and incubated on ice with Fixable Viability Dye eFluor™ 450 (eBioscience) for 20 min. The intracellular staining was carried out according to the manufacturer’s instructions using Fixation/Permeabilization kit (eBioscience) in the presence or absence of antibodies against OCT3/4 (OCT4) (AlexaFluor 647 mouse anti-OCT3/4, BD Biosciences), SOX17 (APC Goat anti-SOX17, R&D system), AFP (AlexaFluor 647 mouse anti-AFP, R&D system) and HNF4A (Alexa Fluor 488 Mouse anti-HNF4A, SantaCruz). For the membrane staining, antibodies against HLA-ABC (FITC Mouse Anti-HLA-ABC, BD Biosciences) and HLA-DR (APC Mouse Anti-HLA-DR, BD Biosciences) were used. The flow cytometry (FACS) analysis was performed with a FACSCanto II (BD Biosciences) and Flow Jo software (Tree star, Ashland, OR, USA).

### Enzyme-linked immunosorbent assay (ELISA) analysis

The quantity of human AFP and HGF secreted into the cell culture medium were determined using the AFP Human ELISA Kit (Abnova) and the HGF Human ELISA Kit (Invitrogen) following manufacturer’s instructions. Human AFP secreted into the sera of animals were determined by the Human AFP Elisa Quantification Kit (Invitrogen, EHAFP) following manufacturer’s instructions.

### Immunofluorescence Assay

Cultured cells were fixed with 4% paraformaldehyde for 15 min at room temperature, permeabilized with 0.5% Triton X-100 in PBS for 15 min and blocked with 1% BSA-0.1% Triton in PBS for 30 min. Primary antibodies were diluted in blocking solution and incubated for 1 hour at room temperature. Secondary antibodies were diluted in blocking solution and incubated for 1 hour at room temperature (primary and secondary antibody information in supplementary Table 2). Cells were mounted using coverslips and ProLong Gold Antifade Mountant (Life Technologies) All pictures were observed under a Zeiss fluorescent microscope.

### Biochemical analysis

Liver injury was estimated by the increase in the serum activities of Alanine-Amino-Transferase (ALAT) and Aspartate-Amino-Transferase (ASAT) measured by the biochemicals laboratory of University Hospital Center (Nantes, France).

### Detection of human Alu DNA sequence by PCR

Genomic DNA was extracted from liver using the Genomic DNA from organs and cells Kit (Macherey-Nagel) following manufacturer’s recommendations. Alu PCR was conducted using two primers: hAluR: 5’-TTT TTT GAG ACG GAG TCT CGC TC-3’ (SEQ ID NO: 1) and hAluF: 5’-GGC GCG GTG GCT CAC G-3’ (SEQ ID NO: 2). PCR was carried out with Herculase Kit (Agilent) in a total volume of 25 μL with 10 ng of genomic DNA. PCR cycling parameters were taken from manufacturer’s instructions. PCR products were run on microfluidic 5K chip in capillary electrophoresis device (Caliper Labchip-GX, LifeScience).

### Histological analysis

Liver tissues were fixed in 4% paraformaldehyde and embedded in paraffin. Sections of 4 μm were used for Hematoxylin-Eosin-Safran (HES) and immunohistochemistry (IHC) staining. After dewaxing, the paraffin slices were incubated 20 min with EDTA buffer (10 mM; pH 9.0) or citrate buffer (10 mM; pH 6.0) at 98°C and endogenous peroxidases were blocked 10 min using 3% H2O2. The presence of human cells was analysed by specific detection of the human Ku80 protein in mouse livers. For this, slices were incubated 45 min in Blocking Solution (Vector Laboratories), then with primary rabbit monoclonal anti-human Ku80 antibody (Abcam) for 1 hour and finally with EnVision+ Dual Link system-HRP (DAB+) (Agilent Dako) for 30 min at room temperature. For Ki67 staining, slices were sequentially incubated in avidin-biotin blocking solution (Invitrogen) for 10 min, in Animal-Free Blocker (Vector Laboratories) for 45 min and with primary rabbit polyclonal anti-mouse/human Ki67 antibody (Abcam) overnight at 4°C. Afterwards, Rabbit Specific HRP/DAB (ABC) Detection IHC Kit (Abcam) was used according to the manufacturer’s manual. All IHC stainings were counterstained with Mayer’s hematoxylin (Sigma-Aldrich). After dehydration, slices were embedded in NeoMount Medium (Merck Millipore) and scanned with a digital slide scanner (Nanozoomer, Hamamatsu Photonics). Files were analysed with the NDP viewer 2.5 software (Hamamatsu).

### Proteomics

An unbiased quantification of all detectable peptides and proteins was performed in liver tissue of no APAP, APAP only and APAP with treatment mice by HRMTM mass spectrometry (Biognosys). Differentially regulated proteins in each cluster/group were identified using the following criteria: q-value <0.025 and average fold change >1.5. Principal component analysis was conducted in R using prcomp and a modified ggbiplot function for plotting, and partial least squares discriminant analysis was performed using mixOMICS package. Functional analysis was performed using g:Profiler and String-db (string-db.org) (Szklarczyk et al., 2019).

### Statistical analysis

Statistics was achieved with the GraphPad Prism5 software. All results were expressed as mean ± SEM. Mann-Whitney, Kruskal-Wallis, One- or two-way ANOVA analysis were used when appropriate. Animal survival was analysed by means of the log rank (Mantel-Cox) test.

## Results

### Differentiation of GStemHep using a cGMP-compliant protocol

We modified our previously protocol (Fourrier et al., 2020) to develop a fully cGMP-compliant production procedures for both human PSC maintenance/amplification and their hepatic differentiation under feeder-free conditions and using only small molecules, recombinant factors and GMP-available reagents and cell culture media. At the end of hepatic differentiation process, cell supernatants were kept for the measurement of the secreted molecules and GStemHep were frozen for the various RNA and protein analyses. The cell identity and gene expression profile were first analysed by RT-qPCR (Figure 1A). The pluripotency-associated markers OCT4 and NANOG were lost in GStemHep while the endoderm-associated genes FOXA2 and SOX17 and the hepatic-associated genes HNF4A and AFP were detected. We could also detect the expression of HGF in GStemHep. Detected gene expression was validated at the protein level by flow cytometry analyses of harvested and frozen cells (Figure 1B). At day 0 of hepatic differentiation, PSC highly expressed OCT4 (> 98%) whereas they did not express anymore OCT4 after differentiation into GStemHep (below the threshold of flow cytometry detection of 0.2%). The average expression of SOX17, HNF4A and AFP markers in GStemHep were 22.3% (± 4.8), 36.2% (± 6.5) and 32.5% (± 7.2), respectively (Figure 1C). All these results were corroborated by immunofluorescence assay with a complete loss of expression of the pluripotency OCT4 and gain of hepatic-specified gene markers SOX17, HNF4A and AFP in GStemHep while PSC were highly positive for OCT4 and negative for AFP and HNF4A (Figure 1D). Interestingly, GStemHep maintained some proliferative activity, as assayed by immunostaining of the cell cycle associated marker Ki67 while as expected, PSC are highly proliferative (Figure 1D). GStemHep secreted AFP and HGF in the cell culture supernatants at the level of 11.2 μg/10^6^ cells and 3.3 ng/10^6^ cells during the last 24h of differentiation, respectively (Figure 1E). As expected, no AFP and HGF was detected in undifferentiated PSC cell supernatant. GStemHep does not express mature hepatocyte-specific markers, such as albumin and CYP3A4 cytochrome CYP450 (not shown). In addition, GStemHep did not show expression of MHC class II (HLA-DR molecule) and lost the expression of MHC class I (HLA-ABC molecule, 2.3% ± 0.6), which is initially detected in PSC (92.4%), as assessed by flow cytometry (Supplementary Figure S1).

**Figure 1.**
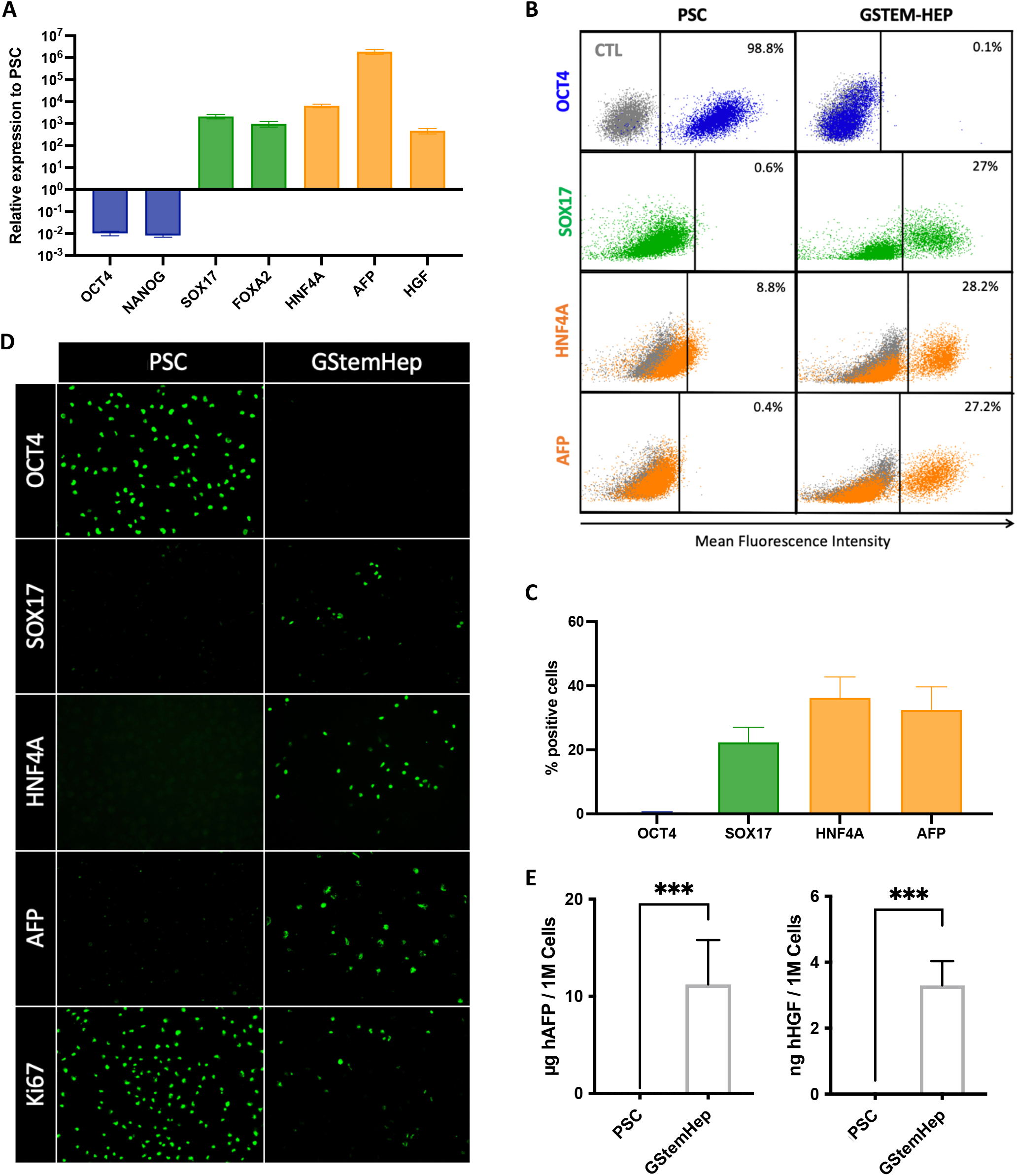
Expression profile of differentiation markers by GStemHep. Human PSC were differentiated into GStemHep with a 10-day protocol. The expression profile of the indicated markers was analysed on cells and culture supernatants. Pluripotency, endoderm and hepatic markers are shown in blue, green and orange, respectively (except for immunofluorescence staining). (A) RT-qPCR analysis of the expression of key markers for differentiation stages and commitment to liver fate in GStemHep (n=10). Results were normalized to GAPDH housekeeping gene and expressed as fold change relative to PSC. (B) Representative FACS plot of a PSC and a GStemHep production batch (detection threshold of OCT4 flow cytometry is 0.2%). Cells incubated with isotype control are in grey. (C) Average expression of pluripotency factor (OCT4) and hepatoblast markers (SOX17/HNF4A/AFP) by GStemHep (n=9). (D) Immunofluorescence staining of GStemHep and human PSC for pluripotent (OCT4), proliferation (Ki67), endoderm (SOX17) and hepatic markers (HNF4A et AFP). (E) Detection of secreted human AFP and human HGF in culture supernatant by ELISA (n=10) (*** p=0.0007 Mann-Whitney test).

In conclusion, we developed a fully compatible process to produce GStemHep cells that have the phenotype of immature hepatic progenitor cells (AFP+/ALB-) secreting human AFP and HGF with some proliferative activity.

### Cryopreservation of GStemHep

To use GStemHep for cell therapy as frozen cells, we evaluated several cGMP-compliant cryopreservation solutions. Here, GStemHep were cryopreserved in CryoStor® CS10 cell freezing medium and stored at −150°C. The cells were then thawed to analyse the impact of freezing on their viability and their replating ability after different time periods of cryopreservation. Viability of harvested and freshly-produced GStemHep is 94.7% ± 0.5 (average of 18 production batches) (Figure 2A). The cell viability was not changed in our cryopreservation and storage conditions overtime, for up to 36 months, as shown in the Figure 2A. Viability of frozen cells was 83.5% (± 5.4) after 36 months of storage, which is not significantly changed as compared to 89.9% (± 0.5) at 0-3 month storage. We showed that cryopreserved GStemHep were able to attach onto culture dishes after thawing, showing that they retain their functional adhesive properties to the recombinant laminin-extracellular matrix for up 12 months (Figure 2B). Maintenance of the immature hepatocyte phenotype of cryopreserved GStemHep was also evaluated by FACS and RT-qPCR analysis. In both analyses, no difference was observed between fresh and thawed cells on expression of pluripotency (loss), endoderm or hepatic markers (gain) (Figure 2C and 2D).

**Figure 2.**
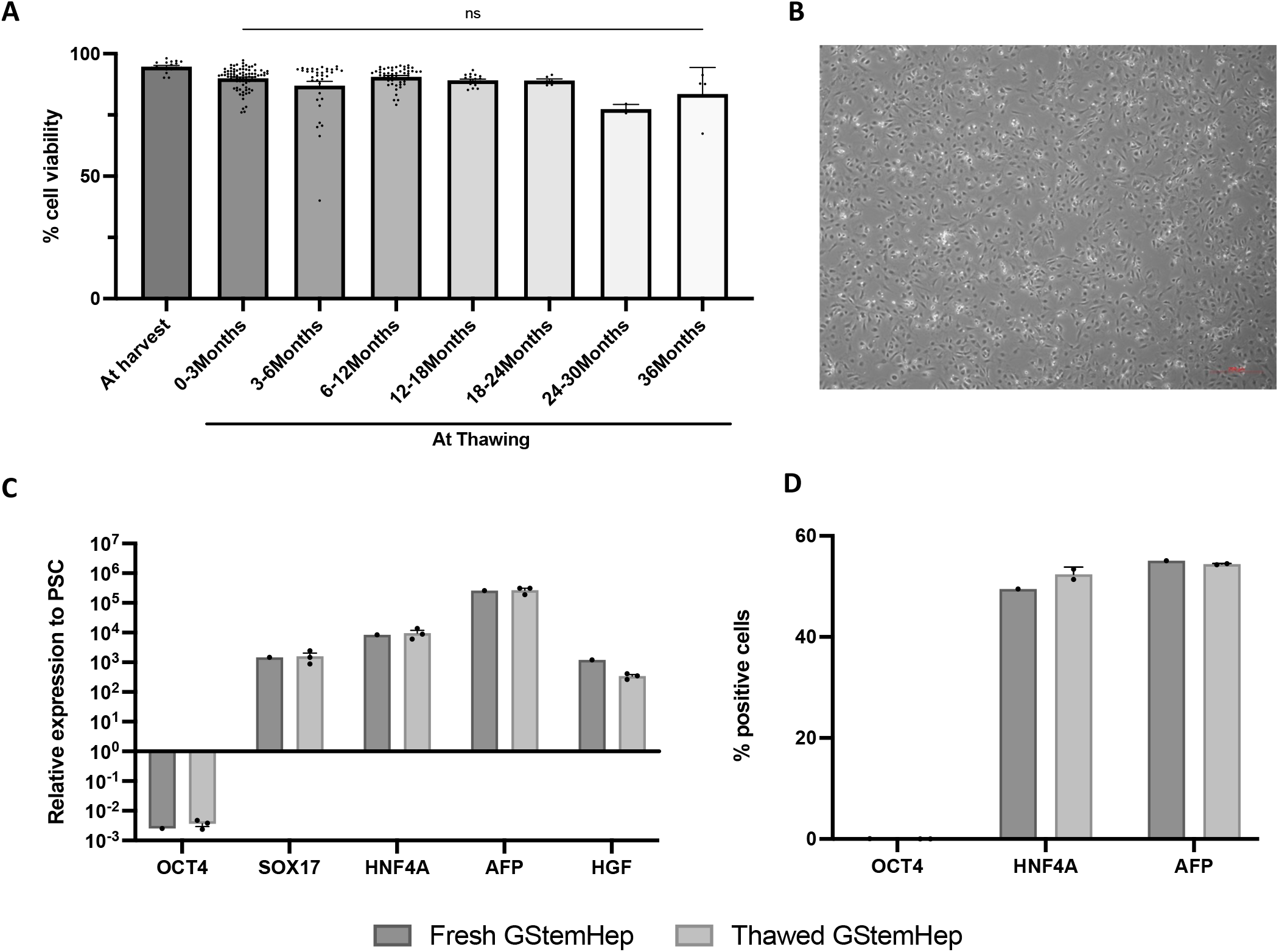
Cryopreservation of GStemHep. After 10 days of differentiation, GStemHep were harvested and frozen in a cryopreservation solution. Cell viability, adhesion capacity and hepatic phenotype were analysed after thawing. (A) Viability of freshly-produced GStemHep at harvest and at thawing after different months of freezing at −150°C; each dot represents a different batch or thawing (n=202) (Kruskal-Wallis test, ns: no significant). (B) Morphology of GStemHep (1 year frozen) 24 hours after thawing and seeding onto a culture plate (Magnification 5x). (C) RT-qPCR and (D) FACS analysis of key marker expression in a cell production batch before and after freezing.

To conclude, the overall freezing conditions of GStemHep were optimized for a high viability after thawing with the maintenance of the cell phenotype and the adhesion capacities, equivalent to that of the freshly-produced cells for a long period storage.

### Frozen GStemHep rescue APAP-induced acute liver failure in immunodeficient mice

To investigate the therapeutic potential of frozen GStemHep for acute liver failure, we used the model of ALF in immunodeficient mice lethally intoxicated with APAP, as previously described (Tolosa et al., 2015). After APAP overdose, 80% of animals died within 4 days with >50% lethality occurring within 2 days (Figure 3A). By contrast, after a unique dose of frozen GStemHep (1×10^6^ cells) that were thawed and injected in the liver via the spleen, most of ALF mice survived, which is statistically different from untreated control mice (p<0.05) (Figure 3A). Noteworthy, this therapeutic benefit was obtained with different cell production cell batches (n=5), cryopreserved for up to 5 months and in different animal studies (n=8, total of 46 treated mice). This increase in survival is accompanied by a significant decrease in the markers of liver damage ASAT and ALAT in the serum of the treated animals at 24h post-injection (Figure 3B). Noteworthy, this decrease was also observed at 3h and 6h after GStemHep injection (Supplementary Figure S2A).

**Figure 3.**
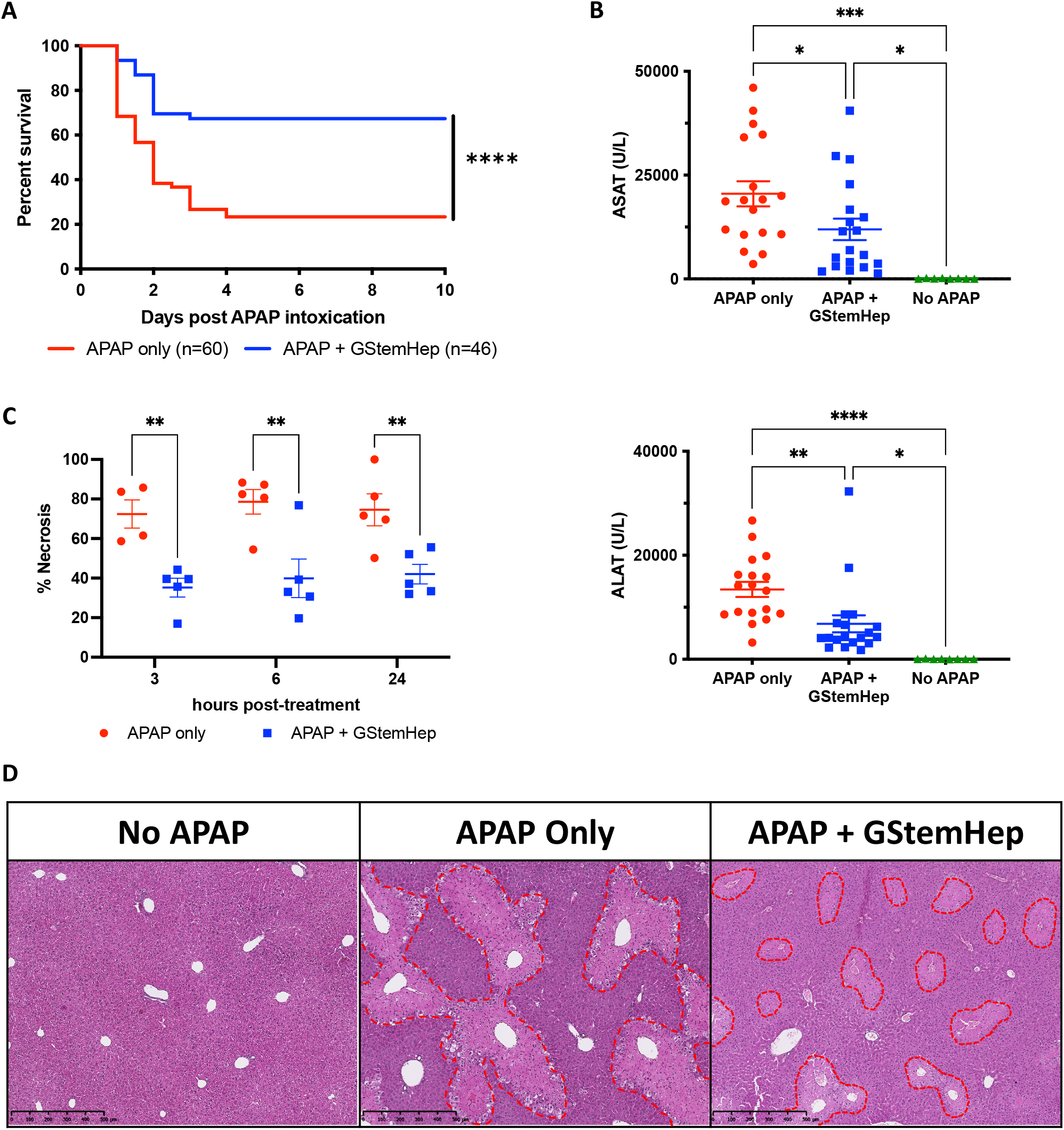
Therapeutic effect of GStemHep in NOD/SCID mice with APAP-induced acute liver failure. After APAP intoxication, mice were treated (APAP+GStemHep) or not (APAP Only) with intrasplenic injection of 1×10^6^ thawed GStemHep. Mice survival was followed over 10 days or mice were euthanized at 3h, 6h and 24h post treatment for analysis of liver damage markers. (A) Survival curve of APAP-induced ALF mice treated with GStemHep injection compared with non-treated group (****p<0.0001 Log-rank (Mantel-Cox) test). (B) Biochemical analysis of liver damage markers: ASAT and ALAT in serum of each group, 24h after APAP intoxication (* p<0.05; ** p<0.005; ***p<0.0005; ****p<0.0001 One-way ANOVA test). (C) Necrosis quantification on HES-stained liver sections in APAP only and APAP + GStemHep groups at 3h, 6h and 24h after GStemHep transplantation (** p<0.01 Two-way ANOVA test). (D) Representative HES-stained sections of liver of each group (n=5 per group), 24h after GStemHep transplantation (Magnification 5x). Areas of necrosis are delimited by the red dotted lines.

The cytoprotective effect of GStemHep was also demonstrated by histopathological comparison of liver tissue. Massive necrosis (>70%) is present in the liver as early as 3h after intoxication, i.e. at the time of GStemHep injection (Supplementary Figure S2B). At 3h post-cell injection, there was a significant decrease in liver necrosis areas (72.5% ±7.1 to 35.3% ±4.8, p<0.05) (Figure 3C). Three hours later i.e., 6h post-cell injection, this reduction is maintained with a necrosis of 40% (±9.8) in treated mice against 78.7% (±6.2) in the APAP control group. At twenty-four hours after intoxication, massive liver necrosis was still histologically observed in the liver of untreated control ALF animals (Figure 3D, middle photo) in contrast to GStemHep-treated ALF animals (Figure 3D, right photo). As expected, no necrosis was observed in the liver of healthy animals that did not received APAP (Figure 3D, left photo). Seven days after the intoxication the necrotic areas are no longer present in the liver of the treated mice (Supplementary Figure S2C).

So, frozen GStemHep rescue mice from a massive and rapid death induced by APAP-intoxication and massive hepatocyte death with a protective effect observed within 24h and as soon as 3h post-cell transplantation.

### GStemHep tracking in the APAP-induced acute liver failure

To demonstrate the presence of GStemHep in the liver, we performed a targeted amplification of human DNA in animal samples. Targeted PCR amplification of human *ALU* sequence demonstrated the specific presence of cells in liver of GStemHep-treated mice 24h after transplantation (Figure 4A). Human *ALU*-PCR analyses were negative at 7 days post transplantation (Supplementary Figure S3A).

**Figure 4.**
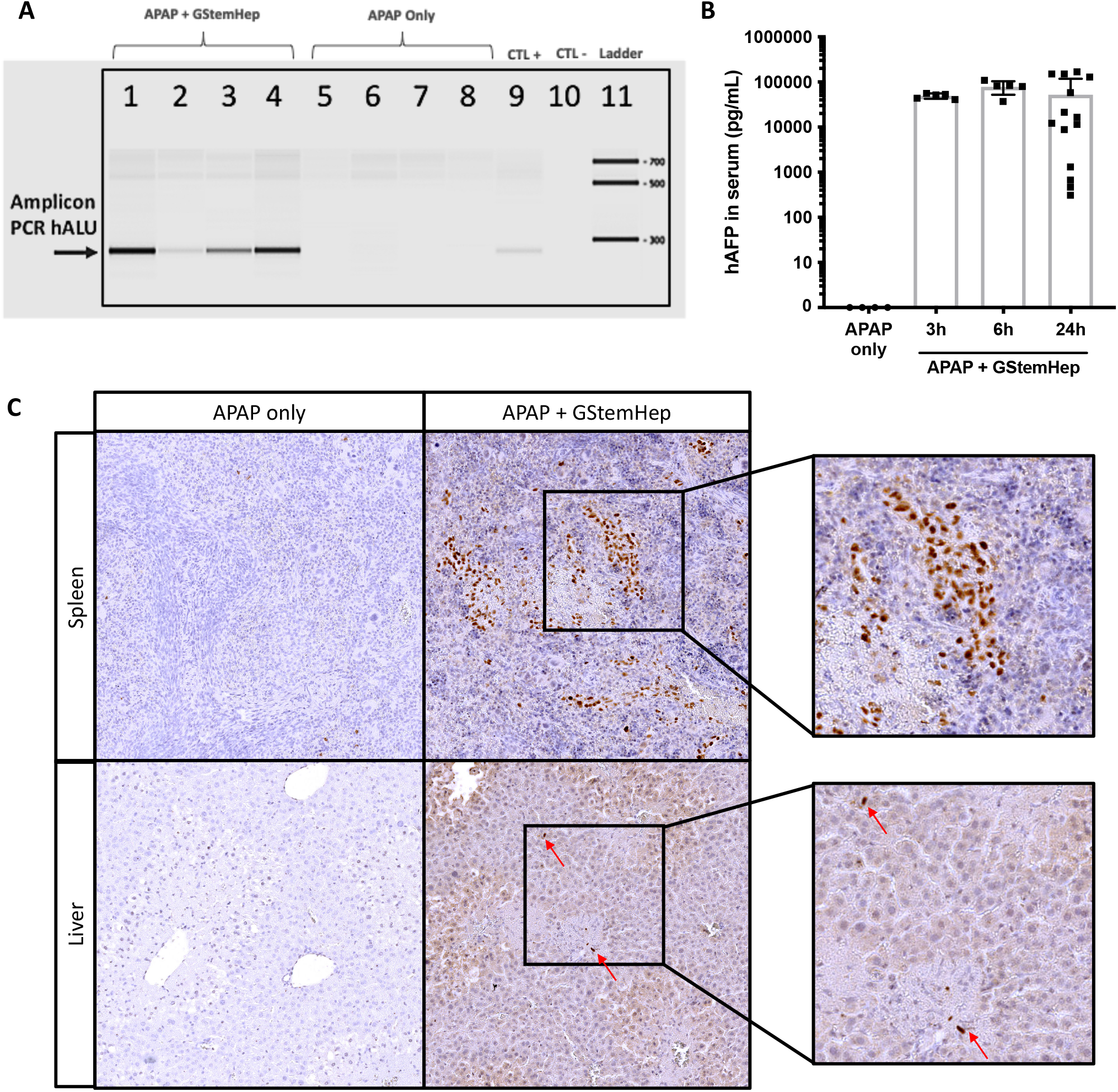
GStemHep tracking in the APAP-ALF model after transplantation. After APAP intoxication and cell transplantation (APAP+GStemHep), cell homing was analysed in the liver and the spleen and human AFP was quantified in serum of mice at different time post-treatment (3h, 6h and 24h). Untreated APAP-ALF mice served as controls (APAP Only). (A) Detection of human ALU DNA sequences (human specific/290bp) in mice liver by PCR at 24h after transplantation, each number represents a different mouse. (B) Quantification of human AFP in mice serum by ELISA 3h, 6h and 24h after transplantation (n=4 in APAP only, n=5 in APAP + GStemHep at 6h and 9h, n=14 in APAP + GStemHep at 24h posttransplantation), each point represents a different mouse. (C) Immunohistochemistry staining for specific human Ku80 in mice liver and spleen at 24h after transplantation (Magnification 10x on the left and 20x on the right), data representative of 10 analysed mice.

We confirmed that GStemHep homed to the livers using human Ku80 immunohistochemistry staining of the liver of GStemHep-treated mice at 24h post-transplantation (Figure 4C). GStemHep were also detected in the spleen, as expected due to the intrasplenic injection. Most of the cells are detected in the spleen at this timing.

Human AFP was detected in mice serum at 3h, 6h and 24h after cell transplantation, confirming that the thawed GStemHep cells were rapidly functional *in vivo* upon thawing and administration (Figure 4B). The mean level of human AFP in mice serum was not significantly different between the 3 time points and ranged from 49.1 ng/mL (±2.9) at 3h post-cell transplantation) to 78.3 ng/mL (±11.6) at 6h post-transplantation. Nevertheless, the quantity of human AFP at 24h post-cell transplantation was more heterogeneous between to the mice.

### GStemHep lead to decreased inflammation and restoration of liver functions

To better understand the therapeutic mechanisms of GStemHep in APAP-induced ALF model, the livers of APAP mice with or without treatment were collected at 3h, 6h and 24h post cell transplantation and analysed compared to healthy mice liver. The inflammation and proliferation liver profile were first analysed by RT-qPCR (Figure 5A). The interleukin-1 receptor antagonist (IL1RN gene), a potent anti-inflammatory antagonist of IL-1 family cytokines which are key inflammatory cytokines in ALF development (Barbier et al., 2019), is statistically increased with the cells in the short term (3h and 6h) as compared to untreated ALF mice. The overexpression of the pro-inflammatory IL-6 cytokine, another hallmark of ALF, was reduced at 3h and 6h in the GStemHep-treated ALF mice compared to untreated ALF mice. Of note, the IL-6 level was still significantly detected in the treated ALF mice above the basal level of control mice with no APAP. The expression of the proliferation Ki67 marker did not differ between the treated or untreated groups. This observation was also confirmed by immunohistochemical labeling of Ki67 on liver tissues (Supplementary Figure S4). Interestingly, gene expression of the vascular endothelial growth factor A (VEGFa) was significantly increased in GStemHep-treated groups at 3h and 6h post-cell transplantation. Twenty-four hours after transplantation, TGFß1 is statistically decreased in animals treated with GStemHep.

**Figure 5.**
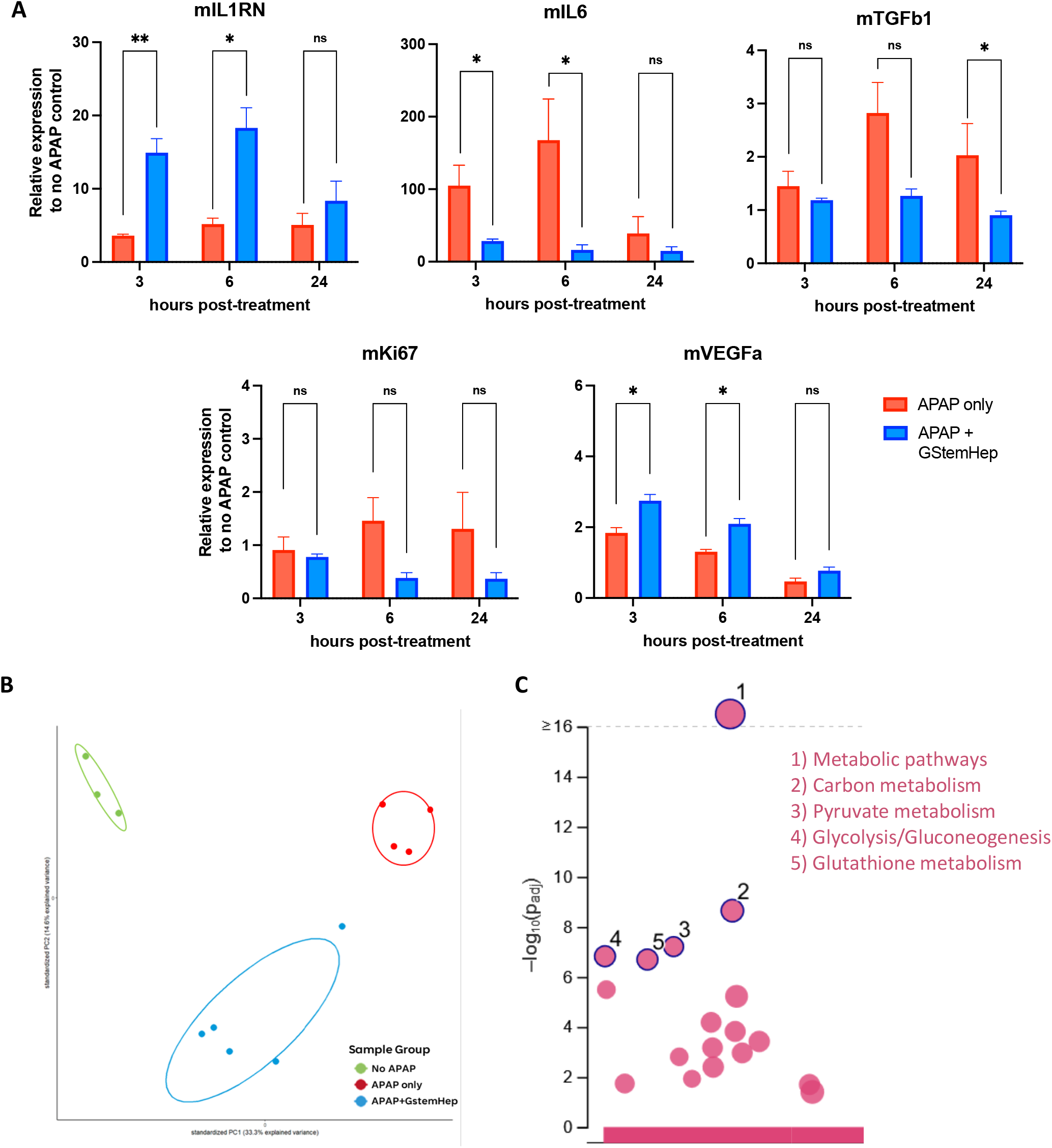
Mechanistic effects of GStemHep in APAP-ALF NOD/SCID mice. After APAP intoxication, mice were treated (APAP+GStemHep) or not (APAP only) with 1×10^6^ thawed GStemHep. Mice livers were collected at 3h, 6h and 24h post cell-transplantation to analyse variations in gene and protein expression. (A) RT-qPCR analysis of expression of inflammation or regeneration markers (mIL1RN, mIL6, mTGFß, mKi67, mVEGFa) in APAP only and APAP+GStemHep groups (n=5 per group). Results were normalized to mGAPDH housekeeping gene and expressed as fold change relative to healthy control mice (* p<0.05; ** p<0.01; ns: no significant, Mann-Whitney test). (B) Proteome-wide data set variance described by Principal Component Analysis (PCA) on healthy (no APAP, n=3), APAP only (n=4) and APAP + GStemHep groups (n=5), 6h after cell transplantation (i.e. 9h post-APAP intoxication). (C) Analysis of differentially-upregulated proteins in APAP+ GStemHep group compared to APAP only group on KEGG pathway.

The therapeutic effect of GStemHep was then evaluated by proteomic analysis using HRM™ mass spectrometry of liver samples from GStemHep-treated ALF mice at 6h-post cell transplantation (APAP+GStemHep), untreated control ALF mice at time corresponding to 6h post-GStemHep cell transplantation (APAP only) and healthy control mice (no APAP). Principal component analysis (PCA) was performed to visualize data set variance in dependence on sample groups (Figure 5B). PCA showed clear separation between the three mice groups, with the main variance PC1 and the secondary variance PC2, explained by difference between the sample groups and by the effect of treatment, respectively. The number of differentially-regulated proteins between healthy mice group and untreated ALF mice group was 861. By contrast, this number was lower between healthy mice group and GStemHep-treated ALF mice group with 275 differentially-regulated proteins, showing that treated ALF mice were closer to healthy mice as compared to untreated ALF mice. There were 320 differentially-regulated proteins between GStemHep-treated and untreated ALF mice groups. KEGG pathway analysis of this differentially-regulated proteins showed 18 KEGG pathways enriched in treated group (Figure 5C). Metabolic pathways were the most up-regulated, including carbon metabolism, glycolysis/gluconeogenesis, pyruvate and glutathione metabolism.

### Frozen GStemHep rescue APAP-induced acute liver failure in immunocompetent mice

To confirm and further highlight the therapeutic potential of GStemHep for ALF, we used a second well-described ALF mice model in immunocompetent C57Bl/6 animals lethally intoxicated with the TAA hepatotoxin. Twenty-four hours after TAA-intoxication, animals received an intrasplenic injection of thawed GStemHep in absence of immunosuppression drug. Immediately after cell thawing and without any cell wash step, GStemHep were injected in the portal vein via intrasplenic administration route. We observed, as for the APAP model, a significant large survival of transplanted mice (69%) compared to untreated mice (Figure 6A). This was demonstrated using 6 different cell production batches (cryopreserved between 2 and 9 months) in 20 different animal studies showing the robustness of therapeutic efficacy of GStemHep, as well as robustness of cell production process. Interestingly, 83% of untreated ALF-control mice died within 48h after transplantation, showing that cell therapy needed to be efficient within the short term to rescue mice from ALF-induced death (Figure 6A).

**Figure 6.**
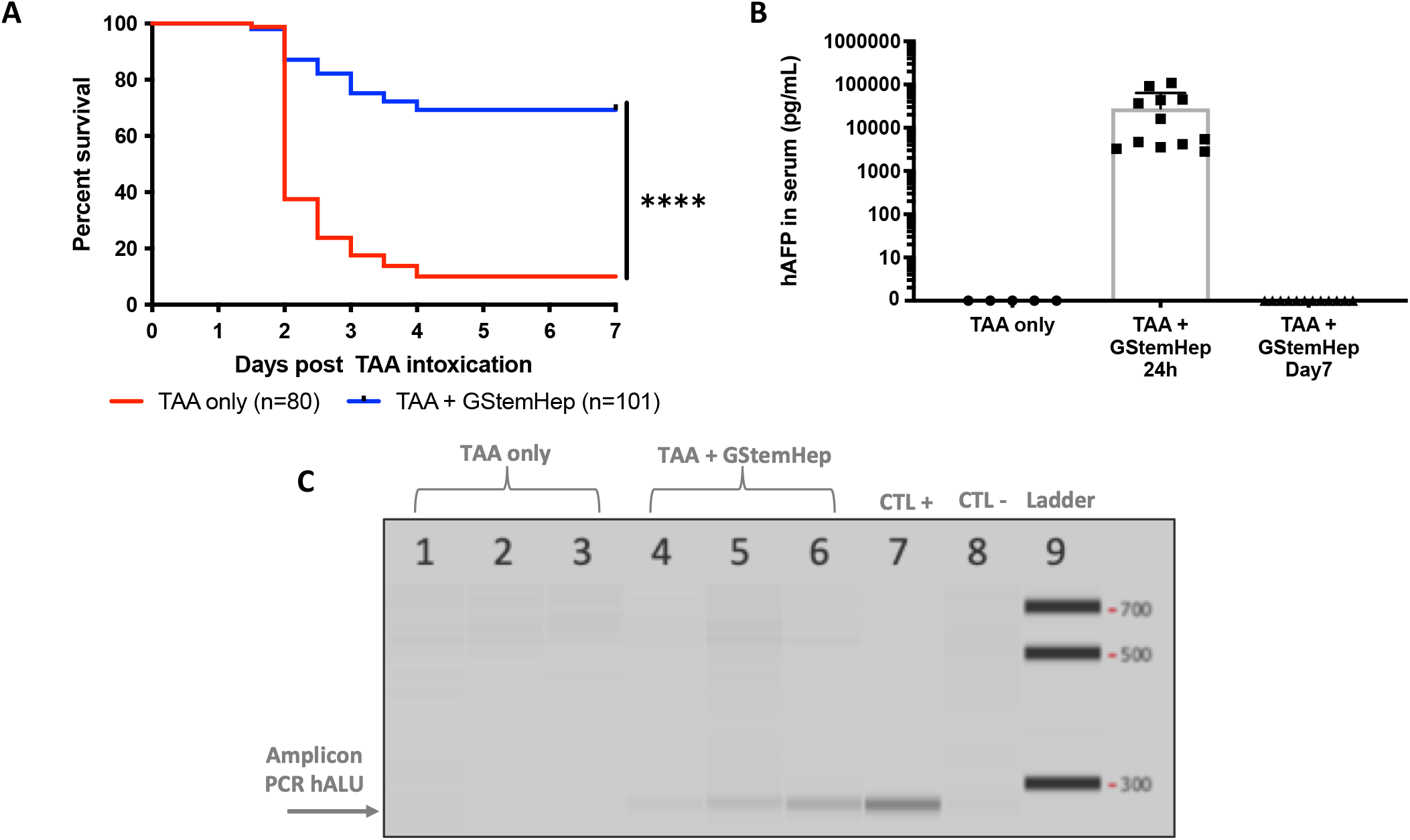
Therapeutic effect of GStemHep and cell tracking in C57Bl/6 mice with TAA-induced ALF. After TAA intoxication, mice were treated (TAA+GStemHep) or not (TAA only) with intrasplenic injection of 1×10^6^ GStemHep. Mice survival was monitored for 7 days, or mice were euthanized at 24h post treatment for PCR analysis in the liver. Sera were collected at different times. (A) Survival curve of TAA-induced ALF mice treated with GStemHep compared with non-treated group (****p<0.0001 Log-rank (Mantel-Cox) test). (B) Quantification of human AFP in serum of TAA only group (n=5) and of TAA+GStemHep group at 24 hours (n=13) and 7 days (n=5) after GStemHep transplantation by ELISA. (C) Detection of human ALU DNA sequences in mice liver by PCR at 24 hours after transplantation, each number represents a different mouse.

Human AFP could be detected in mice serum 24h (28±10 ng/mL on average). Noteworthy, human AFP in mice serum was not detected in after 7 days (Figure 6B). Human *ALU* PCR analyses showed the presence of human DNA in the mice livers 24h after cell transplantation (Figure 6C) but not at 9 days (Supplementary Figure S3B).

## Discussion

The encouraging results of hepatocyte transplantation and the emergence of PSC as an unlimited cell source have paved the way for a new cell therapy of ALF. The challenges of this therapy are the production of hepatic cells, cultured and differentiated in a cGMP compliant manner and on a large scale while keeping therapeutic potency. In view of the emergency, it is also essential to guarantee the availability of the treatment at any time in the clinical context.

In this study, we investigated the potential therapeutic effect of frozen human PSC-derived hepatic progenitors called GStemHep, produced in cGMP compliant feeder-free and serum-free conditions. Indeed, at present, very few proposed protocols are fully cGMP, i.e. for both PSC amplification and hepatocyte differentiation steps (Blackford et al., 2019). Several cGMP-compliant hepatic differentiation protocols were described but PSCs were generally amplified using non-cGMP matrices for PSC culture such as Matrigel (Chen et al., 2020; Fourrier et al., 2020; Tolosa et al., 2015). Here, all raw materials, media, cytokines and matrix used for PSC amplification and differentiation are compatible with cGMP standards.

In addition, all the previous studies were focused on the therapeutic potential of PSC-derived HLCs, i.e. similar to fetal and not fully-differentiated hepatocytes (Blackford et al., 2019; Hannan et al., 2013; Luce et al., 2021; Rashidi et al., 2018; Schwartz et al., 2014; Tolosa et al., 2015). The HLCs were characterized mainly by the expression of the fetal marker AFP, the mature hepatic marker ALB and significant detoxification activities (CYP3A4, ammonia detoxification, …). They showed therapeutic potential when transplanted in animal models with various liver failures including ALF (Chen et al., 2015; Nagamoto et al., 2016; Pettinato et al., 2016; Rashidi et al., 2018; Takebe et al., 2017; Tolosa et al., 2015). In our study, we demonstrate that immature hepatic progenitors (AFP+/ALB-) with no detoxification activities at time of transplantation show therapeutic effects for treating ALF. Noteworthy, the duration of the differentiation protocol of GStemHep is relatively shorten (10 days) as compared to that of HLC (>20 days), resulting in cost and time savings. Interestingly, GStemHep became lowly immunogenic after hepatic differentiation (loss of expression of MHC class I and no expression of MHC class II).

The other major limitation of PSC-based therapeutic treatment is the need of sufficient quantities of cells available at all times. Cell cryopreservation allows to be immediately at disposable at any time and particularly for an emergency use. However, for now, all the various studies showing a therapeutic effect of HLC in animals modeling ALF used freshly-prepared cells (Asgari et al., 2013; Chen et al., 2020; Liu et al., 2011; Takayama et al., 2017; Tolosa et al., 2015). Indeed, cryopreservation and thawing procedures have been reported to have detrimental effects on the viability and function of human hepatocytes when compared to freshly-isolated cells (Stéphenne et al., 2007; Terry et al., 2010). Improved hepatocyte cryopreservation protocols have been described but therapeutic efficacy has not been reported in ALF, yet (Jitraruch et al., 2017; Takayama et al., 2017). We previously reported that frozen PSC-derived hepatic stem cells were able to rescue mice modeling Crigler-Najjar inherited disease, i.e. in a context of a non-fulminant liver diseases where transplanted cells have time to differentiate into fully-mature hepatocytes (Fourrier et al., 2020). The question that remains is whether frozen cells can save liver failure outside of inherited metabolic diseases. Here, the overall freezing conditions of GStemHep were optimized for a high viability after thawing with maintenance of the phenotype and the adhesion capacities of freshly-produced cells. Furthermore, frozen GStemHep displayed a significant therapeutic potential after thawing and immediate transplantation without prior cell washing steps in two ALF mice models. Their resistance to cryopreservation-related damages may be associated to the immature hepatic phenotype of the cells.

The therapeutic effect resulted in a significant increase in survival of ALF mice and a decrease in hepatic injury markers (transaminase, liver necrosis), which was seen as soon as 3h post-cell transplantation in APAP-induced ALF mice model. This showed that the action of the GStemHep is very fast. Moreover, the therapeutic effect was observed in both immunodeficient and immunocompetent models in absence of immunosuppression where xenogeneic immune rejection is stronger. The *in vivo* therapeutic potential was demonstrated in several independent animal studies (8 for APAP-ALF and 20 for TAA-ALF mice) and using different cell batches (up to 6), showing the robustness of therapeutic efficacy of GStemHep, as well as robustness of cell production process. We showed that GStemHep became rapidly functional upon thawing as soon as 3h post-transplantation (presence of human AFP in mice serum) and homed to the liver within 24h (hAlu-PCR, anti-hKu80 immunostaining). GStemHep can be detected as soon as 3h post-transplantation in the liver after splenic administration route and disappears within 7 days. The absence of long-term cell engraftment, the very rapid death of mice as well as the rapidity of the therapeutic effect suggests a paracrine mechanism of action of GStemHep. We demonstrated that GStemHep produced and secreted significant level of HGF, which is the most potent growth factor for hepatocytes (Nagamoto et al., 2016). This secreted HGF probably plays a major role in the therapeutic effects of GStemHep. Indeed, Takayama et al showed the production of HGF by HLCs is key for rescuing mice from ALF (Takayama et al., 2017).

To better understand the mode of action of GStemHep, an unbiased proteomic and gene expression analyses were carried out on animal liver. Due to the rapid observed action of GStemHep, the expression of hepatic inflammation and regeneration genes was analysed 3h, 6h and 24 hours after treatment. Inflammation is reported to play a key role in liver failure and regeneration but this must be finely regulated (Michalopoulos, 2007). Tuning this response could be as important as supporting impaired organ functions. Despite its role in liver regeneration, excess inflammation in ALF leads to more serious damages (Wu et al., 2010). Interestingly, we observed a decrease in IL1RN and IL-6 gene expression as soon as 3h post-cell transplantation and thereafter suggesting that GStemHep may prevent an early excess of inflammation, a hallmark in ALF. For instance, decreasing IL-6-related inflammation has been shown to restore cerebral blood flow and reduce features of hepatic encephalopathy in mice with APAP-induced ALF (Roth et al., 2021) and decrease liver damage (Chowdhury et al., 2019; Coelho et al., 2022). Noteworthy, IL-6 gene expression was still detected in GStemHep-treated mice over the basal level of healthy mice, which may be crucial for promoting liver regeneration of the injured liver. Concomitantly, we observed an increase in VEGFa gene expression at 6 and 24 hours after treatment. As several reports have demonstrated an important role of VEGFa in liver regeneration and hepatocyte proliferation after APAP-induced hepatotoxicity (Donahower et al., 2006; Kato et al., 2011; Papastefanou et al., 2007), an enhanced expression of VEGFa could contribute to the therapeutic effect of GStemHep. No effect on hepatocyte proliferation was observed within 24h post-cell transplantation. With peak of hepatocyte proliferation after major partial-hepatectomy occurring at 48h (Lehmann et al., 2012) and 72h after a high-dose APAP (Bhushan et al., 2014), the analysis of proliferation markers (Ki67) in this study was therefore certainly too early to show a significant increase in liver regeneration after GStemHep treatment.

Unbiased proteomic analyses of liver samples confirmed the different expression profile between treated and untreated APAP-induced ALF mice. The primary analysis of the liver proteome at 6h after cell transplantation also suggests a return towards the basal state of the healthy liver. Indeed, there is a greater expression of the proteins involved in the metabolic pathways of the liver in the treated mice suggesting a more functional liver as compared to untreated ALF mice. Noteworthy, proteomic results at 6h post-GStemHep treatment reveal that TGFß1, which is known to be inhibitory of hepatocyte proliferation after a partial hepatectomy and to inhibit liver regeneration after APAP toxicity, is increased in untreated APAP-ALF mice while significantly decreased in treated APAP-ALF mice to a similar level to that of healthy mice (Bird et al., 2018). These observations are confirmed by the qPCR analysis of TGFß1 gene expression. Concomitantly, we observe a significant increase in expression of WNT/ ßcatenin in GStemHep-treated mice as compared to that of untreated APAP-ALF. It has been shown that an increased expression of WNT/ßcatenin signaling pathway improved liver regeneration after APAP overdose (Bhushan et al., 2014). Deeper data analyses are underway to more precisely define all the molecules and biological pathways that ultimately mediate liver protection from ALF by GStemHep.

In conclusion, we describe the production of GStemHep, which are immature hepatic cells (AFP+/ALB-) secreting HGF and derived from human pluripotent stem cells, using fully cGMP-compliant protocols. We show a clear and rapid therapeutic effect in different models of ALF after injection of a single dose of GStemHep, demonstrating that therapeutic potential is not bound to a mature hepatic phenotype. These GStemHep cells can be cryopreserved and used after thawing without impacting their therapeutic potential. They have a rapid therapeutic effect, as soon as 3h post-cell transplantation, with a decrease in liver injuries and an increase in metabolic functions. Their mode of action relies on alleviation of inhibition factors of liver regeneration, increase in proliferation-promoting factors and decrease liver inflammation. Finally, their therapeutic effects can be demonstrated in immunocompetent animals in absence of immunosuppression. Overall, these results pave the way to conduct a phase I/II clinical trial for a PSC-based regenerative medicine of acute liver failure with a single dose of frozen GStemHep in absence of immunosuppression to repair the liver without graft.

## Acknowledgments

We acknowledge the IBISA MicroPICell facility (Biogenouest), member of the national infrastructure France-Bioimaging supported by the French national research agency (ANR-10-INBS-04) and the cytometry (Laurence Delbos) and molecular biology (Laurent Tesson) platforms of the UMR1064 laboratory.

This work was supported by ANR-14-CE16-0026-StemHepTher, by the European ERDF program (FEDER PAYS DE LA LOIRE n°PL001690), by the RHU program “iLite” on “Innovations for Liver Tissue Engineering” granted by PIA2 through ANR-16-RHUS-0005 and by a fellowship CIFRE-ANRT (2019/0174).

