## Supplementary data for "Cryopreserved cGMP-compliant human pluripotent stem cells-derived immature hepatic progenitors rescue mice from acute liver failure"

**Table S1 : Primers for Taqman assays**

| Gene | Reference |
|---|---|
| **hPOU5F1 (OCT4)** | Hs00999632_g1 |
| **hNANOG** | Hs04260366_g1 |
| **hSOX17** | Hs00751752_s1 |
| **hFOXA2** | Hs00232764_m1 |
| **hHNF4A** | Hs00604435_m1 |
| **hAFP** | Hs00173490_m1 |
| **hHGF** | Hs00300159_m1 |
| **mIL-1RN** | Mm00446186_m1 |
| **mTGFb1** | Mm01178820_m1 |
| **mKi67** | Mm01278617_m1 |
| **mVEGFa** | Mm00437306_m1 |

**Table S2 : Antibodies for Immunofluorescence Assay**

| Protein | Reference | |
|---|---|---|
| **OCT4** | SC9081 | Santa Cruz Biotechnology |
| **SOX17** | AF1924 | R&D Systems |
| **HNF4A** | SC374229 | Santa Cruz Biotechnology |
| **AFP** | A8452 | Sigma-Aldrich |
| **CK19** | M0888 | Dako |
| **Ki67** | Ab15580 | Abcam |
| **Secondary anti-rabbit** | A21206 | Invitrogen |
| **Secondary anti-mouse** | A21202 | Invitrogen |
| **Secondary anti-goat** | A11055 | Invitrogen |

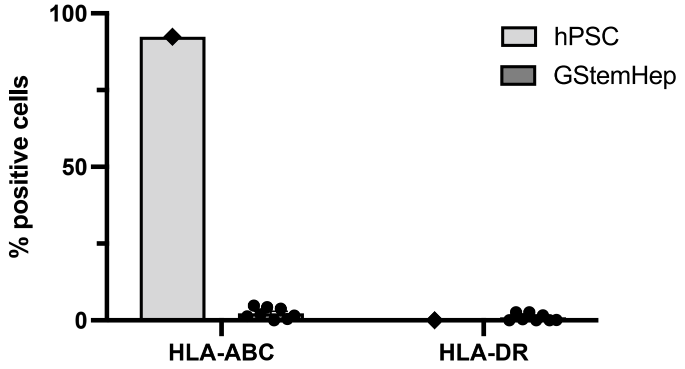


**Figure S1. Immune profile of GStemHep.** Quantification of MHC class I (HLA-ABC) and class II (HLA-DR) molecules on PSC cell line and GStemHep production batches (n=8, each dot represents a cell batch) by flow cytometry.

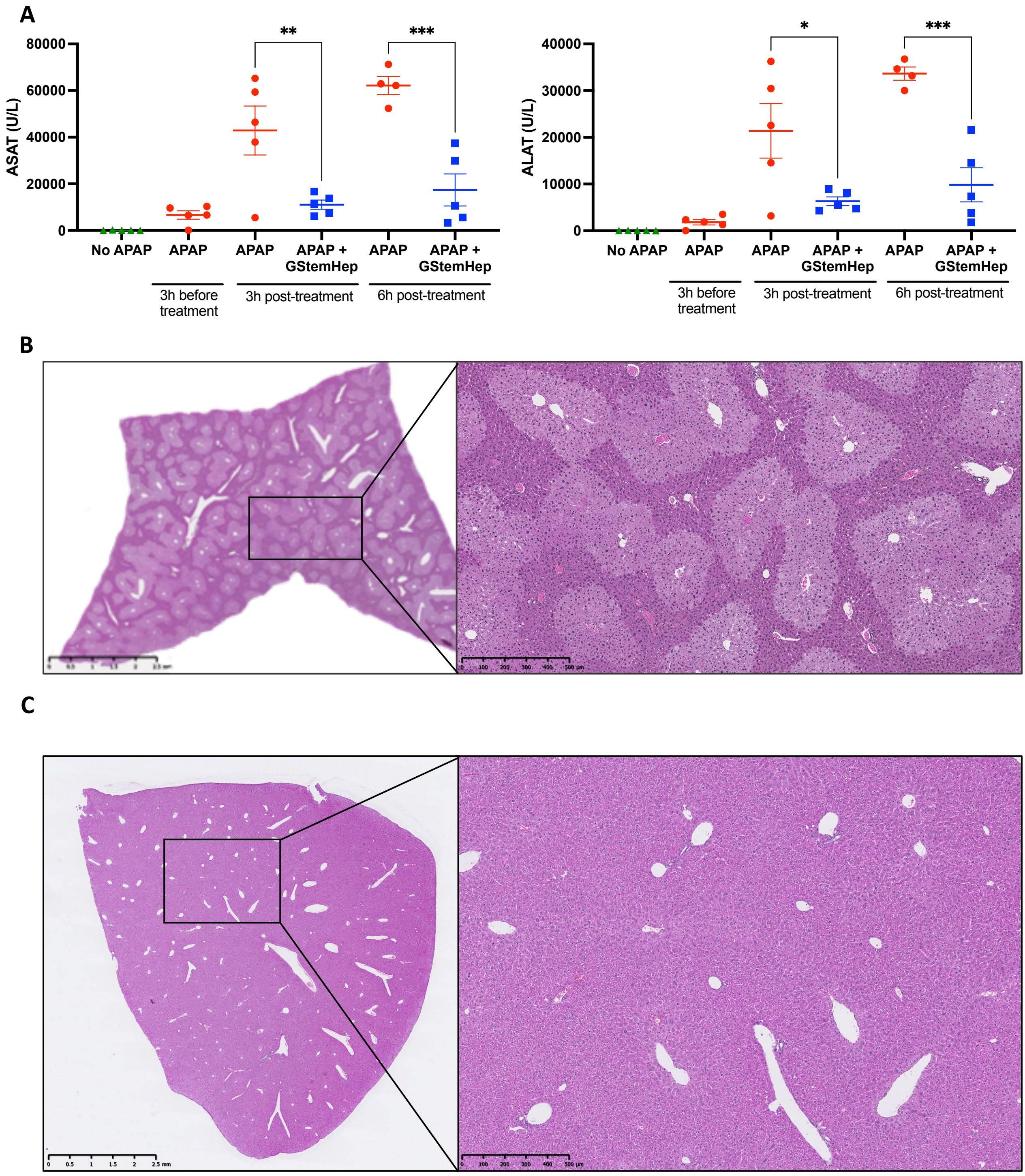


**Figure S2. Therapeutic effects of GStemHep in NOD/SCID mice with APAP-induced ALF.** (A) Biochemical analysis of liver damage markers: ASAT and ALAT in blood serum of each group at 3h and 6h after cell transplantation (* $p<0.05$; ** $p<0.005$; *** $p<0.0005$, One-way ANOVA test). (B) Representative HES-stained sections of liver of at 3h after APAP-injection, i.e. at the time and before GStemHep treatment and (C) 7 days after cell transplantation (Magnification x1 on the left and x5 on the right).

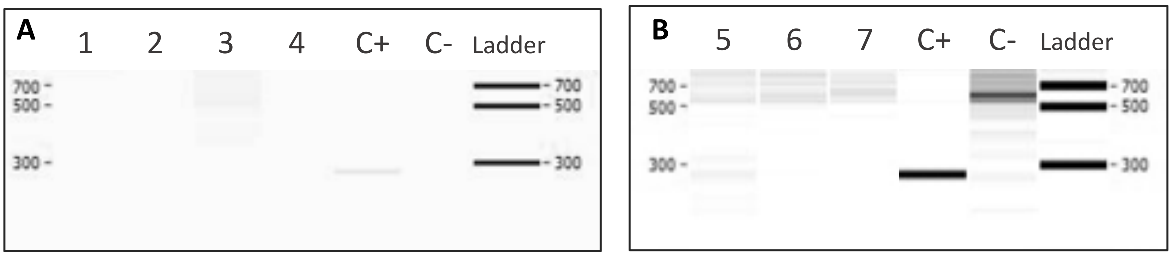


**Figure S3. GStemHep tracking in the APAP- and TAA-induced ALF models at long-term post transplantation.** (A) Detection of ALU DNA sequences (human specific/290bp) in mice liver by PCR at 7 days after GStemHep transplantation in APAP-ALF mice. (B) Detection of ALU DNA sequences in mice liver by PCR at 9 days after transplantation in TAA-ALF mice, each number represents a different mouse (1-4: APAP + GStemHep; 5-7: TAA + GStemHep; C+ : positive control; C- negative control).

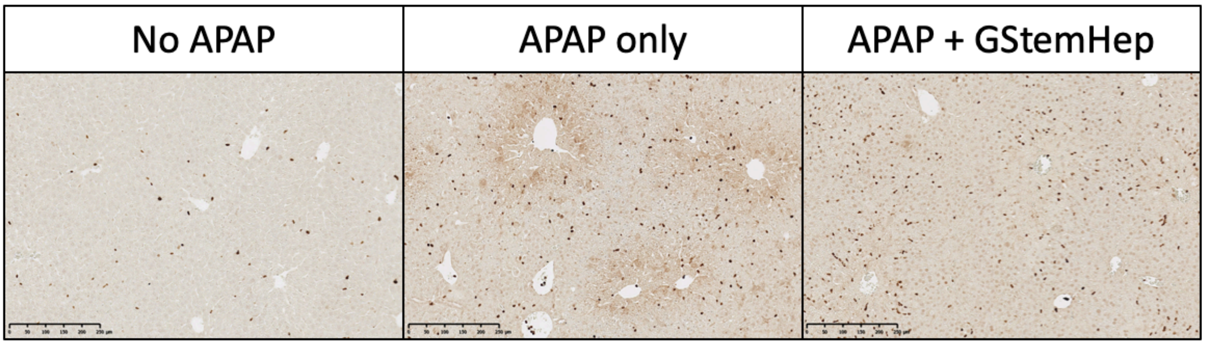


**Figure S4. Evaluation of cell proliferation in the liver of APAP-ALF NOD/SCID mice after GStemHep treatment.** Immunohistochemistry staining for the proliferation marker Ki67 in mice liver of healthy (no APAP), untreated APAP-ALF (APAP only) and GStemHep-treated (APAP + GStemHep) mice at 24h after cell therapy (Magnification 10x), data representative of 5 analysed mice per group.
